# AleRax: A tool for gene and species tree co-estimation and reconciliation under a probabilistic model of gene duplication, transfer, and loss

**DOI:** 10.1101/2023.10.06.561091

**Authors:** Benoit Morel, Tom A. Williams, Alexandros Stamatakis, Gergely J. Szöllősi

## Abstract

**Motivation:** Genomes are a rich source of information on the pattern and process of evolution across biological scales. How best to make use of that information is an active area of research in phylogenetics. Ideally, phylogenetic methods should not only model substitutions along gene trees, which explain differences between homologous gene sequences, but also the processes that generate the gene trees themselves along a shared species tree. To conduct accurate inferences, one needs to account for uncertainty at both levels, that is, in gene trees estimated from inherently short sequences and in their diverse evolutionary histories along a shared species tree.

**Results:** We present AleRax, a software that can infer reconciled gene trees together with a shared species tree using a simple, yet powerful, probabilistic model of gene duplication, transfer, and loss. A key feature of AleRax is its ability to account for uncertainty in the gene tree and its reconciliation by using an efficient approximation to calculate the joint phylogenetic-reconciliation likelihood and sample reconciled gene trees accordingly. Simulations and analyses of empirical data show that AleRax is one order of magnitude faster than competing gene tree inference tools while attaining the same accuracy. It is consistently more robust than species tree inference methods such as SpeciesRax and ASTRAL-Pro 2 under gene tree uncertainty. Finally, AleRax can process multiple gene families in parallel thereby allowing users to compare competing phylogenetic hypotheses and estimate model parameters, such as DTL probabilities for genome-scale datasets with hundreds of taxa

**Availability and Implementation:** GNU GPL at https://github.com/BenoitMorel/AleRax and data are made available at https://cme.h-its.org/exelixis/material/alerax_data.tar.gz.

**Supplementary information:** Supplementary material is available.

## 1 INTRODUCTION

Simultaneously inferring gene trees and the species tree is challenging. Genomes contain abundant information about evolutionary history, but, single gene sequences are often too short to reliably resolve gene trees (Haag *et al*., 2022). Moreover, gene trees are not species trees, but each is the unique result of series of evolutionary events. Starting from an individual site in a genome up to the species level, a hierarchy of evolutionary processes generate genomic sequences, with each level of the hierarchy contributing to the phylogenetic signal that can induce differences between reconstructed gene trees (Szöllősi *et al*., 2015). Segregating mutations that cross speciation events (a process called incomplete lineage sorting) leave topological signatures in gene trees. Duplications, transfers, and losses of genes (DTL) lead to substantial differences in both, the size, and phylogenetic distribution, of families of homologous genes, and at the same time produce patent phylogenetic discord.

While most species tree inference methods take single-copy genes as input that are assumed *a priori* to be orthologous (i.e., to have originated from a common ancestral gene solely through speciation), more recent methods are able to handle more general homologous gene families (i.e., genes that originated from a common ancestor through speciation, gene duplication and possibly transfer events) including multi-copy ones (Boussau *et al*., 2013; Zhang and Mirarab, 2022; Morel *et al*., 2022). In particular, we recently developed SpeciesRax (Morel *et al*., 2022), a tool that models gene duplication, transfer, and loss (DTL) events and that estimates a maximum likelihood (ML) rooted species tree under the so-called UndatedDTL model. Another promising recent method is ASTRAL-Pro 2 (Zhang and Mirarab, 2022), which infers a species tree from multi-copy gene trees by maximizing a duplication-aware quartet score. However, SpeciesRax and ASTRAL-Pro 2 only consider a single *estimated* gene tree per gene family as input, and are thus sensitive to gene tree uncertainty. Alternative methods such as Phyldog (Boussau *et al*., 2013) jointly estimate the species tree *and* the gene trees that evolved along it via speciation, duplication, and loss. However, Phyldog does not scale to large datasets and does not model gene transfer.

Another challenge is to infer a reconciled gene tree, a gene tree together with a reconciliation, a series of DTL and speciation events that trace its evolution along the species tree. To this end, we previously developed GeneRax (Morel *et al*., 2020), a probabilistic method that searches for the reconciled gene trees that maximize the joint likelihood, defined as the product of the phylogenetic likelihood (the probability of the multiple sequence alignment (MSA) given the gene tree) and the reconciliation likelihood (the probability of the gene tree given the species tree under UndateDTL). However, GeneRax does not provide a confidence measure for the inferred gene trees. Amalgamated Likelihood Estimation (ALE) (Szöllősi *et al*., 2013) is an alternative two-step approach that sums the joint likelihood over all reconciled gene trees that can be amalgamated as a combination of clades observed in a sample of gene trees previously inferred from the sequences using conditional clade probabilities (Höhna and Drummond, 2012; Larget, 2013) to approximate the phylogenetic likelihood. Confidence in a particular DTL event can then be estimated by counting how frequently it appears in the different reconciled gene trees. However, ALE is not able to simultaneously process multiple gene families, or to infer the species tree.

Here we present AleRax, a novel probabilistic method for phylogenetic tree inference that can perform both species tree inference *and* reconciled gene tree inference from a sample of gene trees. We first present the method and the key features of AleRax. Then, we present the results of our benchmarks: we found that AleRax is on par with ALE in terms of gene tree reconstruction accuracy, but an order of magnitude faster and robust to numerical underflow that causes ALE to fail. Finally AleRax is 25% more accurate than SpeciesRax and twice as accurate as ASTRAL-Pro 2 for species tree inference.

## 2 METHOD

### 2.1 Likelihood definition and computation

Let *S* be the species tree, and *A* be an alignment of homologous genes, that is, a gene family alignment. The probability of observing *A* given *S* is obtained by summing over all possible rooted gene trees *G* for alignment *A*:

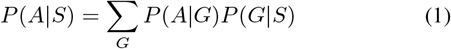

(Felsenstein, 1988; Szöllősi *et al*., 2013).

AleRax approximates *P* (*A* | *S*) by using the ALE (Szöllősi *et al*., 2013) algorithm, which takes as input a sample of gene trees for each gene family, ideally computed via a Bayesian inference tool such as Phylobayes (Lartillot *et al*., 2013) or MrBayes (Ronquist *et al*., 2012) prior to running AleRax. The gene trees in the samples can be rooted or unrooted. Instead of naïvely computing the sum ∑_*G*_ *P* (*A*| *G*)*P* (*G* | *S*), AleRax uses the sample of gene trees provided to estimate a nonzero probability for all gene tree topologies that can be “amalgamated” (David and Alm, 2011) from clades present in the sample using conditional clade probabilities (CCPs) (Höhna and Drummond, 2012; Larget, 2013). This corresponds to the maximum entropy distribution given marginal split frequencies in the sample of gene trees provided as input (Szöllősi *et al*., 2013). Using CCPs allows AleRax to exploit a joint dynamic programming recursion that efficiently and accurately approximates Equation 1 (Szöllősi *et al*., 2013). An in-depth description of this algorithm is provided in the Supplementary Material. Importantly, unlike ALE, AleRax uses arbitrary-precision floating point values when necessary, allowing it to process datasets that cause ALE to fail due to numerical underflow. AleRax can also process multiple gene families in parallel given a sample of genes for each family allowing it to infer the species tree as well as more realistic DTL models, e.g., branch-wise DTL parameters on the species tree, by maximizing the likelihood of the species tree, which can be written as the product over all per-family alignment probabilities:

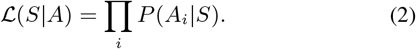

### 2.2 Species tree inference

AleRax searches for the rooted ML species tree. It implements the same tree search strategy as SpeciesRax (Morel *et al*., 2022), which applies hill-climbing subtree prune and regraft (SPR) moves until it cannot find a move that yields a tree with a better likelihood. However, unlike SpeciesRax, which relies on a single gene tree per gene family, AleRax uses a sample of per family gene trees to integrate over gene tree uncertainty by approximating *P* (*A*_*i*_ |*S*). AleRax recomputes the model parameters (described in the next subsection) after each round of SPR moves. The starting species tree can be generated at random, estimated with MiniNJ (Morel *et al*., 2022) (a duplication-aware distance method), or specified by the user.

### 2.3 Model parameter estimation

In general, the reconciliation model used by AleRax, the so called UndatedDTL model (Morel *et al*., 2020) can have three free parameters per gene family *k* and per species tree branch 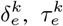, and 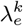, representing the duplication, transfer, and loss probabilities, respectively. However, estimating such a large number of free parameters (proportional to the product of the number of species in *S* multiplied by the number of genes families) can lead to over-parameterization. Hence, AleRax implements three distinct approaches to parameter estimation that can be chosen by the user. In the *global parameters* mode, all families and all species share the same set of parameters *{δ, τ, λ}*, resulting in only three free parameters. In the *per-family parameters* mode, AleRax optimizes a different set of parameters *{δ*_*k*_, *τ*_*k*_, *λ*_*k*_*}* for each gene family *k*. This corresponds to 3*K* free parameters, where *K* is the number of gene families. In the *per-species parameter* mode, the user can provide a list of species groupings that share the same set of parameters. In this mode, the model has 3*n* free parameters, where *n* is the number of groupings defined. In each mode, AleRax optimizes the model parameter values with respect to the likelihood via gradient descent.

### 2.4 Sampling reconciled gene trees

AleRax samples, per gene family, *r* reconciled gene trees proportional to their joint likelihood using stochastic backtracking following the dynamic programming recursion (Szöllősi *et al*., 2013) used to approximate Equation 1. We describe the algorithm in the Supplementary Material. The number of gene trees *r* to sample is specified by the user. For each gene family, AleRax outputs the *r* reconciled gene trees, the rooted majority-rule consensus gene tree, and the number of gene events per species, averaged over all output samples for this specific gene family. The reconciliations are stored in RecPhyloXML format (Duchemin *et al*., 2018) and can be converted into SVG figures using Thirdkind (Penel *et al*., 2022), as illustrated in Figure 1. AleRax also outputs the species pairs involved in horizontal gene transfers, sorted by the number of times those transfers have been sampled (per family and summed over all families). In addition, for each species, it returns the number of ancestral gene copies and the number of DTL events. We also provide scripts to extract lists of families that were involved in transfers between specific species pairs, or families that experienced a specific event for a given species (e.g., all families with a gene duplication in *Arabidopsis thaliana*).

**Fig. 1:**
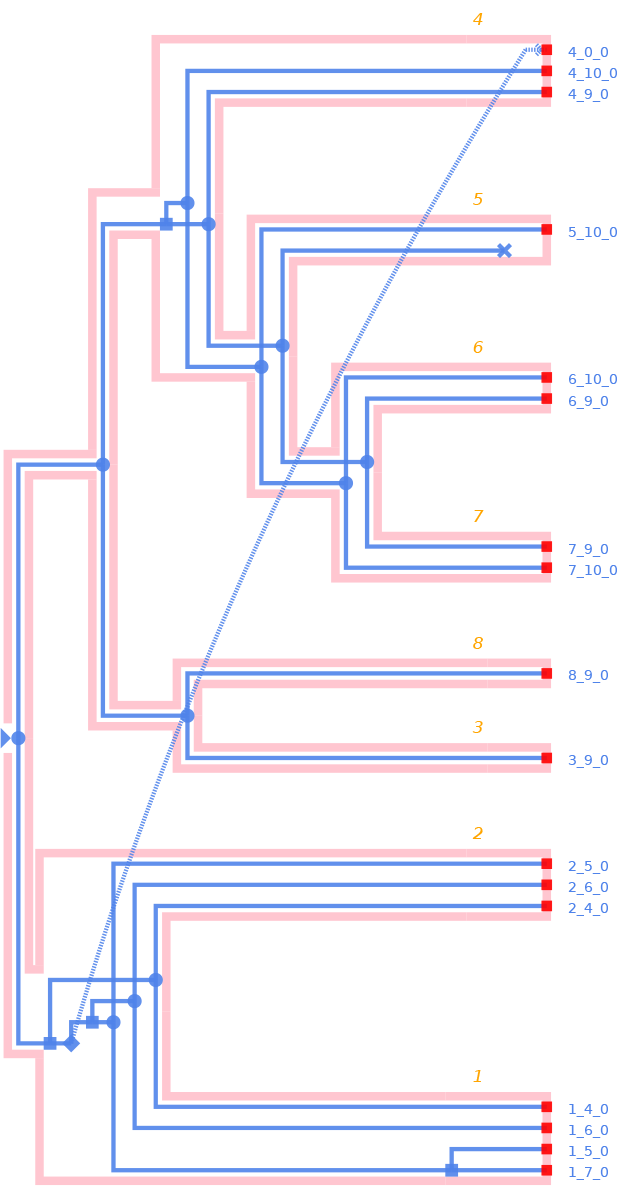
One reconciled gene tree sampled with AleRax and visualized with Thirdkind. The orange labels correspond to species labels, and the blue labels to gene lables.

## 3 RESULTS

Here we only provide a summary of the experiments and their results. We describe them in detail in the Supplementary Material.

### 3.1 Gene tree reconciliation

We ran both ALE and AleRax on the simulated datasets used to benchmark SpeciesRax (Morel *et al*., 2022). As expected, ALE and AleRax perform analogously in terms of accuracy (Fig. 2). We also executed both tools on a real dataset derived from the HOGENOM (Penel *et al*., 2009) database’s HOGENOM-CORE subdataset, which contains 666 representative genomes spanning the diversity of cellular life. ALE failed for 25 families (out of 12, 408) because of numerical underflow. AleRax in contrast successfully processed all families. In terms of runtime, AleRax was on average an order of magnitude faster than ALE, both on simulated and empirical datasets (Fig. 2).

**Fig. 2:**
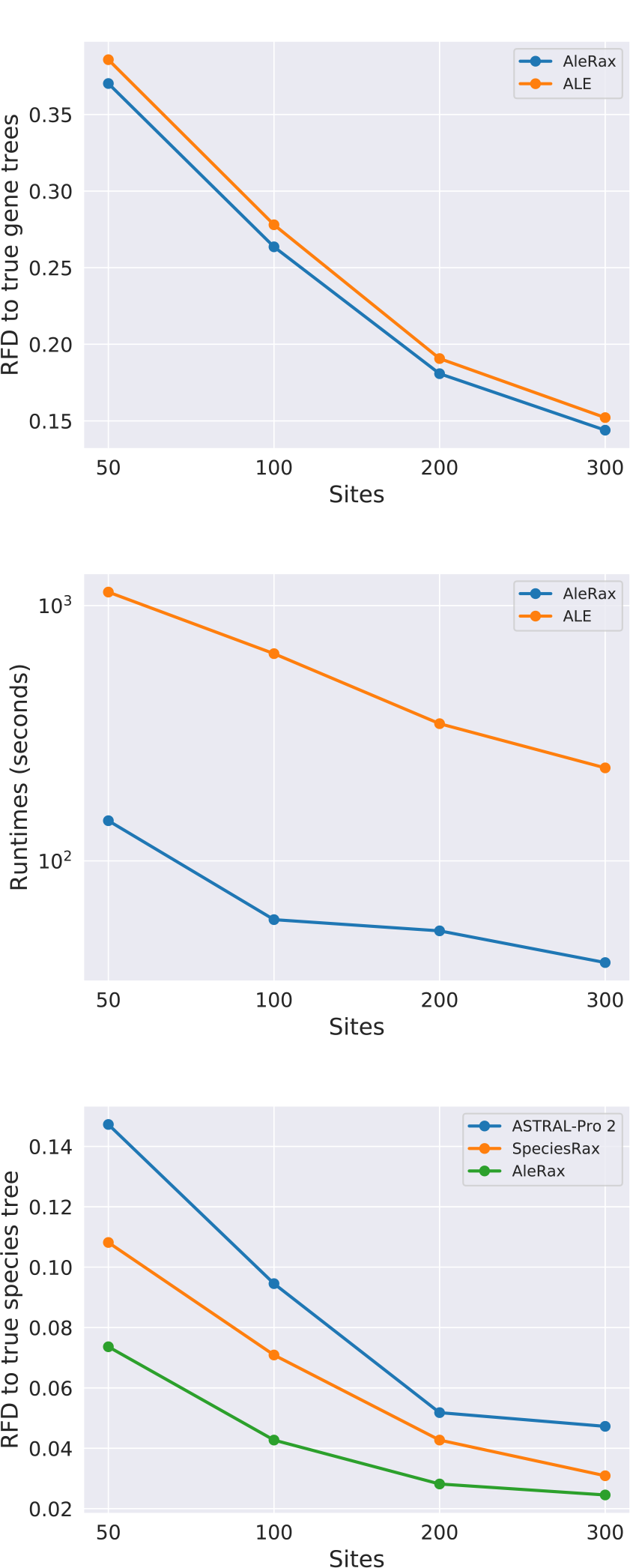
Results on simulated datasets for varying numbers of sites per gene family: gene tree accuracy, runtimes for gene tree inference, and species tree accuracy.

### 3.2 Species tree inference

We investigated the impact of using gene tree distributions instead of single ML gene trees by comparing AleRax, SpeciesRax, and ASTRAL-Pro 2 on simulated datasets. We used the same benchmarks that were employed for assessing SpeciesRax (Morel *et al*., 2022), that is, we varied different simulation parameters individually while keeping the rest fixed, and measured the distance between the true and the inferred species trees. We ran AleRax on gene tree distributions inferred with MrBayes (Ronquist *et al*., 2012). We found that, over all conditions tested, AleRax is on average 25% more accurate than SpeciesRax and twice as accurate as ASTRAL-Pro 2. For instance, Fig. 2 compares the accuracy of the different methods when varying the number of sites per gene family. This accuracy improvement is due to the ability of AleRax to accommodate phylogenetic uncertainty.

## 4 CONCLUSION AND FUTURE WORK

AleRax is an efficient and user-friendly tool for species tree inference and gene tree-species tree reconciliation that can be applied to datasets from across the tree of life. It is particularly useful in biological applications where not only the species tree, but also the histories individual gene families are of interest. We showed that AleRax is on par with ALE in terms of reconciled gene tree accuracy, while being one order of magnitude faster and more robust to numerical errors. Furthermore, it infers more accurate species trees than SpeciesRax and ASTRAL-Pro 2, because it can accommodate gene tree uncertainty. In the future, we plan to improve our model to accommodate DTL rate heterogeneity over species, incomplete lineage sorting, and horizontal gene transfer time constraints.

## ACKNOWLEDGEMENT

BM and AS are financially supported by the Klaus Tschira Foundation, by DFG grant STA 860/6-2, and by the European Union (EU) under Grant Agreement No 101087081 (Comp-Biodiv-GR). GSz received funding from the European Research Council under the European Union’s Horizon 2020 research and innovation programme under grant agreement no. 714774 and the grant GINOP-2.3.2.-15-2016-00057. This work was funded by the Gordon and Betty Moore Foundation through grant GBMF9741 to TAW and GSz.

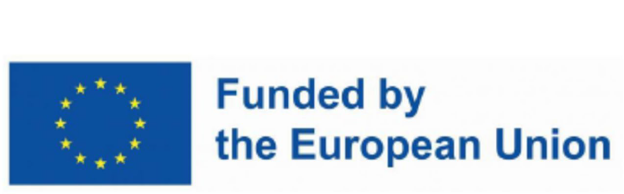

