## Supplementary material for "AleRax: A tool for gene and species tree co-estimation and reconciliation under a probabilistic model of gene duplication, transfer, and loss"

#### 1 Method

In this section we detail the likelihood computation as implemented in AleRax. We first describe the model used to describe gene evolution under gene duplications, transfers, and losses. Then, we explain how Conditional clade probabilitys (CCPs) can be used to approximate the posterior distribution of gene tree topologies. Finally, we derive our algorithm for computing the likelihood of a species tree under the UndatedDTL model using CCPs.

#### 1.1 The UndatedDTL model

The UndatedDTL model, is a discrete state model. The process starts with the **origination** of a single gene copy on branch $e$ of the species tree $S$ with probability $p_e^O$. Subsequently, gene copies evolve independently until, either all copies are observed at the leaves or become extinct. A gene will undergo one of the following events:

- on branch $e$ of the species tree, the gene copy either:
  - **duplicates** with probability $p_e^D$ and is replaced by two descanted genes on the same branch;
  - is **transferred** with probability $p_e^T$ to a random recipient branch that is *not* ancestral to the donor branch, but otherwise drawn uniformly at random from the species tree. In other words, it is replaced by two descanted genes, one on the original donor branch and another on the recipient branch;
  - is **lost** with probability $p_e^L$;
  - or undergoes a **speciation** with probability $p_e^S = 1-(p_e^D - p_e^T - p_e^L)$, as a result of which:
    - on internal branches it is replaced by two descanted genes, one on each descendant branch;
    - on terminal branches it is replaced by a single descendant copy on the subtending leaf.
- on leaf $l$ of the species tree it is **observed** with probability $p_l^{\text{obs.}}$ or, alternately, it is discarded.

Gene trees are rooted trees spanned by observed gene copies with internal nodes corresponding to a speciation, duplication, or transfer event, as illustrated in Fig. 1.

We denote by $\delta_e$, $\lambda_e$, and $\tau_e$ the duplication, loss, and transfer intensity parameters that determine the above event probabilities on branch $e$ of the species tree :

$$p_e^D = \delta_e/(1+\delta_e+\tau_e+\lambda_e) \tag{1}$$
$$p_e^T = \tau_e/(1+\delta_e+\tau_e+\lambda_e) \tag{2}$$
$$p_e^L = \lambda_e/(1+\delta_e+\tau_e+\lambda_e) \tag{3}$$
$$p_e^S = 1/(1+\delta_e+\tau_e+\lambda_e). \tag{4}$$

Note that the parameters $\delta_e$, $\lambda_e$, and $\tau_e$ can be different for distinct families and can conversely be shared for groups of species tree branches. The parameter $p_l^{obs.}$ is the probability of a gene that is present in a genome (leaf $l$ of the species tree $S$) is actually observed in the data set in question. By default $p_l^{obs.} = 1$, but maybe specified per leaf by the user based on, e.g., estimates of genome completeness.

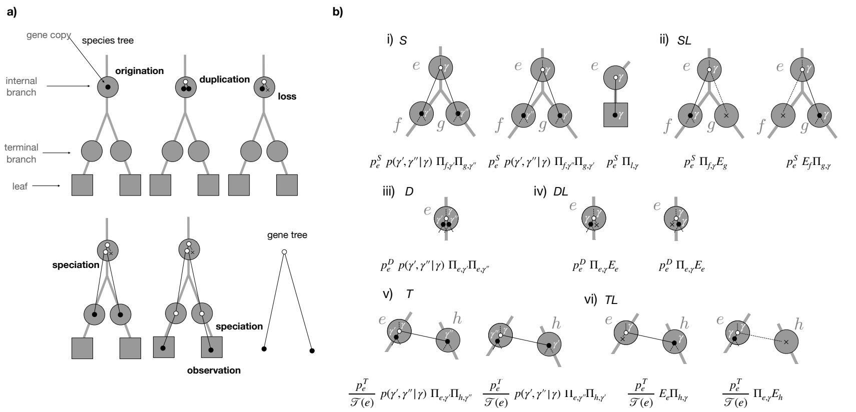


Figure 1: The UndatedDTL model: **a)** a simple scenario, where gene copies (filled black circles) undergo a series events (in bold) along the species tree generating the gene tree. **b)** terms in (18) and (20) required to calculate $\Pi_{\gamma,e}$, that is, the sum over all reconciliations generating the sub-clade $\gamma$ of $\Gamma$ starting from a single gene present on the internal branch $e$ of the species tree $S$. We must consider the following events: i) if $e$ is an internal branch of $S$, speciation with probability $p_e^S$ such that the descendant clade $\gamma'$ of $\gamma$ is observed on $f$ of $S$ and $\gamma''$ on $g$ of $S$, or vice versa. Alternatively, if $e$ is a terminal branch, with probability $p_e^S \Pi_{\gamma,l}$ the clade $\gamma$ will be observed at the leaf $l$ subtending $e$; ii) if $e$ is any branch of $S$, speciation with probability $p_e^S$ such that $\gamma$ is observed on $f$ of $S$ and the copy on $g$ goes extinct with probability $E_g$, or vice versa; iii) duplication with probability $p_e^D$ such that $\gamma'$ and $\gamma''$ are both observed on $e$; iv) duplication with probability $p_e^D$ such that either the first or second copy goes extinct, each with probability $E_e$ and $\gamma$ is observed on $e$; v) transfer with probability $p_e^T$, such that the respective sub-clades $\gamma'$ and $\gamma''$ correspond to the copy on the donor branch $e$ of $S$, while the other copy corresponds to the recipient copy on branch $h$ of $S$ that is not an ancestor of $e$ and finally vi) transfer with probability $p_e^T$ followed by the extinction of either, the copy in the donor linage $e$ with probability $E_e$ of the extinction of the copy in the recipient, with probability $E_h$.

#### 1.2 Conditional clade probabilities

CCPs can be used to accurately approximate the posterior distribution of tree topologies (Larget, 2013) from samples recorded via standard Markov Chain Monte Carlo (MCMC) phylogenetic inference (Höhna and Drummond, 2012). CCPs yield accurate estimates of the posterior gene tree probability for a large number of gene tree topologies, even when the MCMC procedure samples only a minuscule proportion of the overall tree space (Szöllősi *et al.*, 2013).

As described below, the CCP estimates represent the posterior probability of gene trees based on marginal split frequencies, but ignore dependencies between clades. While this independence assumption is generally incorrect, CCP estimates based on sufficiently large samples of trees usually yield very accurate approximations of the posterior probabilities (Höhna and Drummond, 2012). Furthermore, ignoring dependencies between clades is not an arbitrary assumption, because the CCP-based estimate of the posterior probability corresponds to the maximum entropy distribution (Jaynes, 2003) given the marginal split frequencies observed from an MCMC sample of phylogenetic trees (Szöllősi *et al.*, 2013).

From a methodological perspective, CCP-based analyses typically operate on a sample size spanning between 3,000 and 10,000 tree topologies, even in contexts where the actual number of potential tree topologies is considerably greater. In practice, for reasons of computational tractability, several studies have used maximum likelihood, in particular via the IQTree ultrafast-bootstrap mode, instead of MCMC samples.

##### 1.2.1 The probability of a rooted tree topology

We define a clade $\gamma$ as a nonempty subset of a set of homologous sequences $\Gamma$ comprising an alignment $A$. Let $G$ be a rooted tree spanning $\Gamma$, $g$ a subtree of $G$, and $\gamma$ the clade defined by the leaf set of $g$. The probability $q_G(\gamma)$ of the subtree $g$ resolving $\gamma$ in $G$, given CCPs based on a sample of gene trees is given by the recursion:

$$q_G(\gamma) = p(\gamma', \gamma''|\gamma)\ q_G(\gamma')\ q_G(\gamma''),\ \text{if } |\gamma > 1|, \tag{5}$$

where $\gamma'$ and $\gamma''$ are the complementary daughter clades splitting $\gamma$ in $g$, such that $\gamma \setminus \gamma' = \gamma''$ and $p_G(\gamma', \gamma''|\gamma)$ is the *conditional probability* of observing the split ($\gamma'$, $\gamma''$) assuming the clade $\gamma$ is present in $G$.

The conditional probability $p(\gamma', \gamma''|\gamma)$ can be estimated from the marginal frequency of splits in an Markov Chain Monte Carlo (MCMC) sample for clades with at least two genes as:

$$p(\gamma', \gamma''|\gamma) \approx \frac{f(\gamma', \gamma''|\gamma)}{f(\gamma)},\ \text{if } f(\gamma) > 0 \text{ for } |\gamma| > 1, \tag{6}$$

and 0 if $f(\gamma) = 0$, where $f(\gamma', \gamma''|\gamma)$ is the frequency of the split of $\gamma$ into daughter clades $(\gamma', \gamma'')$ and $f(\gamma)$ is the frequency of the mother clade $\gamma$ in the sample of gene trees. The posterior probability of $G$ can be computed recursively, starting from the ubiquitous clade $\Gamma$ containing all leaves of $G$, with the recursion terminating at the single gene clades:

$$q_G(\gamma) = 1, \text{ if } |\gamma| = 1. \tag{7}$$

We refer to gene tree topologies that are entirely composed of clades observed in an MCMC sample of trees as trees that can be *amalgamated* (David and Alm, 2011). As defined here, the CCP estimate of the posterior probability is nonzero for trees that can be amalgamated, and zero otherwise. Note that, the approximated posterior probability is by construction normalized, i.e.:

$$\sum_G q_G(\Gamma) = 1,$$

where the sum is over all rooted tree topologies over the set of leaves $\Gamma$.

If the sample of gene trees is unrooted, which is typically the case in practice due to the time reversibility of most substitution models, each bipartition defines a split $(\Gamma', \Gamma'')$ of the ubiquitous clade $\Gamma$, and for each such split, we set $p(\Gamma', \Gamma''|\Gamma) = \frac{1}{2n-3}$ where $n = |\Gamma|$ is the number of homologous sequences in $A$.

##### 1.2.2 The number of trees that can be amalgamated

To develop intuition for the recursion in the following section utilizing CCPs consider the problem of calculating the number of trees with a nonzero posterior probability given an input sample of gene trees on $n$ leaves, i.e. the number of trees that can be amalgamated.

Denoting the number of rooted trees that can be amalgamated on the complete set of leaves as $N(\Gamma)$ we can recursively calculate their number. To do so we can use the recursion for a clade $\gamma$ with two or more leaves:

$$N(\gamma) = \sum_{\gamma',\gamma''|\gamma} \delta(\gamma', \gamma''|\gamma)\ N(\gamma')\ N(\gamma''), \text{ if } |\gamma| > 1, \tag{8}$$

where the sum $\sum_{\gamma',\gamma''|\gamma}$ is over all splits of the clade $\gamma$ into complimentary subclades $\gamma'$ and $\gamma''$ and $\delta(\gamma', \gamma''|\gamma) = 1$ if the split of $\gamma$ into $\gamma'$ and $\gamma''$ is seen in the sample and zero otherwise. The recursion terminates for clades with a single leaf, i.e., $|\gamma| = 1$, where we have:

$$N(\gamma) = 1, \text{ if } |\gamma| = 1. \tag{9}$$

#### 1.3 Efficiently calculating the joint likelihood

AleRax assumes that genes evolve under the UndatedDTL model. We recently introduced an algorithm to compute $P(G|S)$ for a *single* gene tree (Morel *et al.*, 2020), by summing over all possible sequences of D, T, L, and S events (henceforth called "scenarios") that yield $G$ under the UndatedDTL model. We now adapt this algorithm to approximate the probability of observing a gene alignment $A$ given a species tree $S$ under a given model of gene evolution by summing over all gene trees $G$ using the CCP approximation described above, that is we perform the following calculation:

$$P(A|S) = \sum_G P(A|G) \times P(G|S)$$
$$\approx \sum_G q_G(\Gamma) \times \frac{\sum_e p_e^O P_{e,R}}{\sum_e p_e^O(1-E_e)}, \tag{10}$$

where the sum over $G$ corresponds to the sum over all rooted gene trees, $\Gamma$ is the ubiquitous clade containing all leaves, the sum over $e$ corresponds to the sum over all species tree branches $e$ where the gene family may originate, $p_e^O$ is the probability of origination on branch $e$, and $P_{e,R}$ is the probability of the gene lineage leading to the root $R$ of $G$ being present on branch $e$ as defined in Morel *et al.* (2020). We divide by $\sum_e p_e^O(1-E_e)$ to condition on survival, using the probability of extinction $E_e$ defined below.

Importantly, instead of separately computing each element of the sum in (10), a computationally tractable strategy is to compute for each species branch $e$ and for each gene clade $\gamma$ the probability $\Pi_{e,\gamma}$ of the lineage leading to the first bifurcation resolving the clade $\gamma$ observed on branch $e$ of $S$. Using this approach, introduced in Szöllősi *et al.* (2013), the likelihood can then be expressed as:

$$P(A|S) \approx \sum_G q_G(\Gamma) \times \frac{\sum_e p_e^O P_{e,R}}{\sum_e p_e^O(1-E_e)} = \frac{\sum_e p_e^O \Pi_{e,\Gamma}}{\sum_e p_e^O(1-E_e)}. \tag{11}$$

##### 1.3.1 The extinction probability

We first need to estimate $E_e$, the extinction probability of species $e$, defined as the probability that a gene copy observed on an internal branch $e$ becomes extinct before being observed at the tips of the species tree. We obtain its value by summing over all scenarios that do not yield any observed descendants:

- For an internal branch $e$ of $S$:

$$E_e = p_e^L + p_e^S(E_f E_g) + p_e^D\left(E_e^2\right) + p_e^T\left(E_e \bar{E}_e\right), \tag{12}$$

  and the terms correspond to the i) loss probability, ii) speciation and subsequent extinction probability in both descending lineages (this term must be omitted for terminal branches), iii) duplication and subsequent extinction probability of both copies, and, finally, iv) transfer and subsequent extinction probability of both, the donor copy on branch $e$, and the transferred copy on branch $h$. For the latter event we introduced the notation:

$$\bar{E}_e = \sum_{h \in \mathcal{T}(e)} \frac{E_h}{|\mathcal{T}(e)|}, \tag{13}$$

  where $\mathcal{T}(e)$ is the set of species that can receive an horizontal gene transfer (HGT) from $e$. By default these are all the branches that are *not* ancestors of $e$.

- For a terminal branch $e$ of $S$:

$$E_e = p_e^L + p_e^S E_l + p_e^D \left(E_e^2\right) + p_e^T \left(E_e \bar{E}_e\right), \tag{14}$$

where $l$ is the leaf subtending $e$ in $S$.

- Finally, on leaves $l$ of $S$ the recursion finishes with:

$$E_l = 1 - p_l^{\text{obs.}}, \tag{15}$$

where, as described above $p_l^{obs.}$ is the probability of a gene that is actually present on leaf $l$ of $S$ is observed, consequently $(1-p_l^{\text{obs.}})$ is the probability that a gene not in the dataset was in fact present but was not observed.

In (12) and (14), the value of $E_e$ depends on $\bar{E}_e$, and thus on the extinction probabilities of all species in the species tree. We iteratively estimate $\bar{E}_e$ and $E_e$ for all nodes $e$ in the species tree, by initializing $[E_e]^0 = 0$ and computing for internal branches:

$$\begin{aligned}[E_e]^n =& p_e^L + p_e^S [E_f]^{n-1} [E_g]^{n-1} + p_e^D([E_e]^{n-1})^2 \\ &+ p_e^T [E_e]^{n-1} \sum_{h \in \mathcal{T}(e)} [E_h]^{n-1} / |\mathcal{T}(e)|\end{aligned} \tag{16}$$

and for terminal branches:

$$\begin{aligned}[E_e]^n =& p_e^L + p_e^S E_l + p_e^D([E_e]^{n-1})^2 \\ &+ p_e^T [E_e]^{n-1} \sum_{h \in \mathcal{T}(e)} [E_h]^{n-1} / |\mathcal{T}(e)|,\end{aligned} \tag{17}$$

##### 1.3.2 Dual recursion over reconciliations and gene tree topologies

To carry out the sum over all reconciliations of all amalgamated trees in (10) efficiently we employ a dual recursion over the species tree and over the CCPs that represent the gene trees that can be amalgamated:

- For an internal branch $e$ of $S$:

$$\begin{aligned}\Pi_{e,\gamma} = & p_e^S \sum_{\gamma',\gamma''|\gamma} p(\gamma',\gamma''|\gamma)(\Pi_{f,\gamma'}\Pi_{g,\gamma''} + \Pi_{f,\gamma''}\Pi_{g,\gamma'}) & (S) \\ & + p_e^S \; (\Pi_{f,\gamma} E_g + \Pi_{g,\gamma} E_f) & (SL) \\ & + p_e^D \sum_{\gamma',\gamma''|\gamma} p(\gamma',\gamma''|\gamma)\Pi_{e,\gamma'}\Pi_{e,\gamma''} & (D) \\ & + 2p_e^D \; \Pi_{e,\gamma} E_e & (DL) \\ & + p_e^T \sum_{\gamma',\gamma''|\gamma} p(\gamma',\gamma''|\gamma)(\Pi_{e,\gamma'}\bar{\Pi}_{e,\gamma''} + \Pi_{e,\gamma''}\bar{\Pi}_{e,\gamma'}) & (T) \\ & + p_e^T \; (\Pi_{e,\gamma}\bar{E}_e + \bar{\Pi}_{e,\gamma} E_e) & (TL),\end{aligned} \tag{18}$$

where $f$ and $g$ are the descendants of $e$ in S, and where we have introduced the notation:

$$\bar{\Pi}_{\gamma,e} = \sum_{h\in\mathcal{T}(e)} \frac{\Pi_{\gamma,h}}{|\mathcal{T}(e)|}, \tag{19}$$

where the sum $\sum_{\gamma',\gamma''|\gamma}$ is over all splits of the clade $\gamma$ into complementary subclades $\gamma'$ and $\gamma''$ seen in the sample, abd $\mathcal{T}(e)$ denotes the branches of $S$ to which transfer is allowed from $e$ (by default branches that are *not* ancestors of $e$).

- For a terminal branch $e$ of $S$:

$$\begin{aligned}
\Pi_{e,\gamma} = & \; p_e^S \; \Pi_{l,\gamma} && (S)\\
& + p_e^D \sum_{\gamma',\gamma''|\gamma} p(\gamma',\gamma''|\gamma)\Pi_{e,\gamma'}\Pi_{e,\gamma''} && (D)\\
& + 2p_e^D \; \Pi_{e,\gamma}E_e && (DL)\\
& + p_e^T \sum_{\gamma',\gamma''|\gamma} p(\gamma',\gamma''|\gamma)(\Pi_{e,\gamma'}\bar{\Pi}_{e,\gamma''} + \Pi_{e,\gamma''}\bar{\Pi}_{e,\gamma'}) && (T)\\
& + p_e^T \; (\Pi_{e,\gamma}\bar{E}_e + \bar{\Pi}_{e,\gamma}E_e) && (TL),
\end{aligned} \tag{20}$$

  where $l$ is the leaf subtending terminal branch $e$ in $S$. The terms of (18) and (20) are illustrated in Figure 1b.

- Finally, on every leaf $l$ of $S$, the recursion finishes with:

$$\Pi_{\gamma,l} = \sigma_{\gamma,l} + (1-\sigma_{\gamma,l})(1-p_l^{\text{obs.}}), \tag{21}$$

  where $\sigma_{\gamma,l} = 1$ if $|\gamma| = 1$ and this gene maps to the species tree leaf $l$ and zero otherwise. As described above $p_l^{obs.}$ is the probability of a gene that is actually present being on leaf $l$ of $S$ is observed, consequently $(1-p_l^{\text{obs.}})$ is the probability that a gene not in the dataset, hence not mapping to leaf $l$, was is in fact present but not observed. By default $p_l^{obs.} = 1$, but the values of $p_l^{obs.}$ may be provided by the user, based, e.g., on estimates of genome completeness.

Similar to the expression for the extinction probability $E_e$, the probability $\Pi_{e,\gamma}$ also depends on itself. We solve this through fixed point iteration analogously to (12). Aside of the self dependence, terms only involve some combination of subclades ($\gamma'$ and $\gamma''$) of $\gamma$ or descendants of branch $e$ (branches $f$ and $g$ or a subtending leaf $l$) on $S$. This allows to construct a bottom-up dynamic programming recursion starting at the leaves of $S$ and single gene clades for which $|\gamma| = 1$.

#### 1.4 Sampling reconciled gene trees

AleRax can sample an arbitrary number of reconciled gene trees under the joint likelihood via stochastic backtracking along the sum as follows. We first randomly sample the origination species from the sum in Eq. 11. Then, we recursively sample the sequence of D, T, L, and S events from the sum in (18). Note that the most likely reconciled gene trees can be recovered by replacing the summation in (18) and (20), including in $\sum_{\gamma',\gamma''|\gamma}$, with taking the maximum.

### 2 Simulations

All data generated for this study can be downloaded at https://cme.h-its.org/exelixis/material/alerax_data.tar.gz

#### 2.1 DLSIM and DTLSIM simulations

We generated simulated datasets with SimPhy (Mallo *et al.*, 2015) to assess the influence of the simulation parameters on the reconstruction accuracy of both, the species tree inference (AleRax vs SpeciesRax vs ASTRAL-Pro 2), and reconciled gene tree inference (AleRax vs ALE) methods. We reused the benchmarking setup from (Morel *et al.*, 2022), that we describe again here for the sake of completeness.

The parameters we studied are: the average number of sites per gene family, the number of families, the size of the species tree, the average gene duplication, transfer, and loss (DTL) rates, the gene tree branch length scaler, and the population size. For each parameter studied, we varied its value while keeping all other parameters fixed. In addition, we varied the T rate while keeping the DL rates fixed. We generated 50 replicates for each set of parameter values. We executed the entire experiment twice, once including HGTs (D**T**LSIM experiment) and once excluding HGTs (DLSIM experiment). In the D**T**LSIM experiment, we simulated under the distance-independent HGT model (i.e., the receiving species is uniformly sampled from all contemporary species) and we set the default average HGT rate equal to the default average duplication rates. We provide a detailed list of the SimPhy parameters in Table 1.

#### 2.2 ALESIM simulations

We reused the dataset originally used to benchmark ALE (Szöllősi *et al.*, 2013). Szöllosi *et al.* initially inferred gene trees for 1099 Cyanobacteria gene families using ALE. Then, they simulated new gene sequences under the LG+$\Gamma$+I model along these trees, retaining both, the sequence lengths, and branch lengths.

### 3 Benchmarking species tree inference

In this section, we describe how we compared the species tree inference accuracy of AleRax, SpeciesRax, and ASTRAL-Pro 2 on the DLSIM and DTLSIM simulated datasets.

We inferred the maximum likelihood gene trees with ParGenes (Morel *et al.*, 2018), performing one RAxML-NG search on a single random starting tree per

| Parameter name | Parameter value |
|---|---|
| Standard parameters | |
| Replicates number | 50 |
| Speciation rate | $5 \times 10^{-9}$ |
| Extinction rate | $4.9 \times 10^{-9}$ |
| Number of gene families | 100 |
| Number of species | 25 |
| Dup and loss rates | $\delta \times \textit{Log-}\mathcal{N}(0,1)$, $\delta = 4.9 \times 10^{-10}$ |
| HGT rate | $\tau \times \textit{Log-}\mathcal{N}(0,1)$, $\tau = 4.9 \times 10^{-10}$ |
| GC rate | 0 |
| Population size | 10 |
| Species tree height | $\textit{Log-}\mathcal{N}(21.25, 0.2)$ |
| Global substitution rate | $\textit{Log-}\mathcal{N}(-21.9, 0.1)$ |
| Lineage specific rate gamma shape | $\textit{Log-}\mathcal{N}(1.5, 1)$ |
| Family specific rate gamma shape | $\textit{Log-}\mathcal{N}(1.551533, 0.6931472)$ |
| Gene tree branch specific rate gamma shape | $\textit{Log-}\mathcal{N}(1.5, 1)$ |
| Sequence length | $\nu \times \textit{Log-}\mathcal{N}(0, 0.25)$, $\nu = 100(e^{-\frac{0.25^2}{2}})$ |
| Sequence base frequencies | Dirichlet(A=36,C=26,G=28,T=32) |
| Sequence transition rates | Dirichlet(TC=16,TA=3,TG=5, CA=5,CG=6,AG=15) |
| Seed | $[3000, 3100[$ |
| Varying parameters | |
| Dup and loss rate multiplier | 0.5,1.0,2.0,3.0 |
| HGT rate multiplier | 0.5, 1.0, 2.0, 3.0 |
| Population size | 10, $10^7$, $10^8$, $10^9$ |
| Number of species | 15, 25, 35, 50, 75 |
| Number of gene families | 50, 100, 200, 500, 1000 |
| Average number of sites | 50, 100, 200, 300 |
| gene family tree (GFT) branch length multiplier | 0.01, 0.1, 1, 10.0, 100.0, 1000.0, 10000.0 |

Table 1: SimPhy parameters to simulate the SIMDL and SIMDTL datasets. In the varying parameters section, the rate multipliers are used to scale the constants $\lambda$ for the dup-loss rates and $\tau$ for the HGT rates. For sequence length, $\nu$ is set such as to obtain 100 sites on average.

gene family under the general time reversible model of nucleotide substitution with four discrete gamma rates (GTR+G4) (Tavaré *et al.*, 1986; Yang, 1993). We inferred the distributions of gene trees with MrBayes (Ronquist *et al.*, 2012), with one independent run starting from the maximum likelihood gene trees, two coupled chains, $100,000$ generations, and sampling trees every 100 generations.

Note that, those parameters do not guarantee convergence and might not be optimal for our analyses. We deliberately used a small number of generations to keep the runtimes as well as the $CO_2$ footprint of the entire experiment (the main bottleneck being MrBayes) within reasonable limits. This choice has potential negative impact on *our* new method (AleRax) only when comparing it to SpeciesRax and ASTRAL-Pro 2.

We ran ASTRAL-Pro 2 from the maximum likelihood gene trees using the following command:

```
astral-pro -a mapping.txt -o outputtree.newick -t 40
    raxml_gene_trees.newick
```

We ran SpeciesRax from the maximum likelihood gene trees using the following command:

```
mpiexec -np 40 generax -f families.txt -s MiniNJ --si-strategy
    HYBRID --strategy SKIP  --si-strategy HYBRID
    --per-family-rates --skip-family-filtering
```

We ran AleRax from the gene tree distributions using the command:

```
mpiexec -np 40 alerax -f families.txt -s MiniNJ
    --gene-tree-samples 0 --species-tree-search HYBRID
```

Finally, for each dataset, we assessed the species tree reconstruction accuracy by computing the average relative RF distance between each inferred species tree and the corresponding true species tree using the ETE Toolkit (Huerta-Cepas *et al.*, 2016).

We summarize the results in Figures 2 and 3. AleRax is on average 25% more accurate than SpeciesRax. However, the accuracy of the methods and how they compare to each other substantially depends on the simulation parameters. As expected, the accuracy of both methods decreases with increasing gene tree error: this happens for low numbers of sites and for extreme (short or long) branch lengths. Furthermore, the accuracy decreases when the incongruence between the gene trees and the species trees increases (for instance with high DTL rates or large population sizes). We note that AleRax outperforms SpeciesRax on *all* tested simulation parameters.

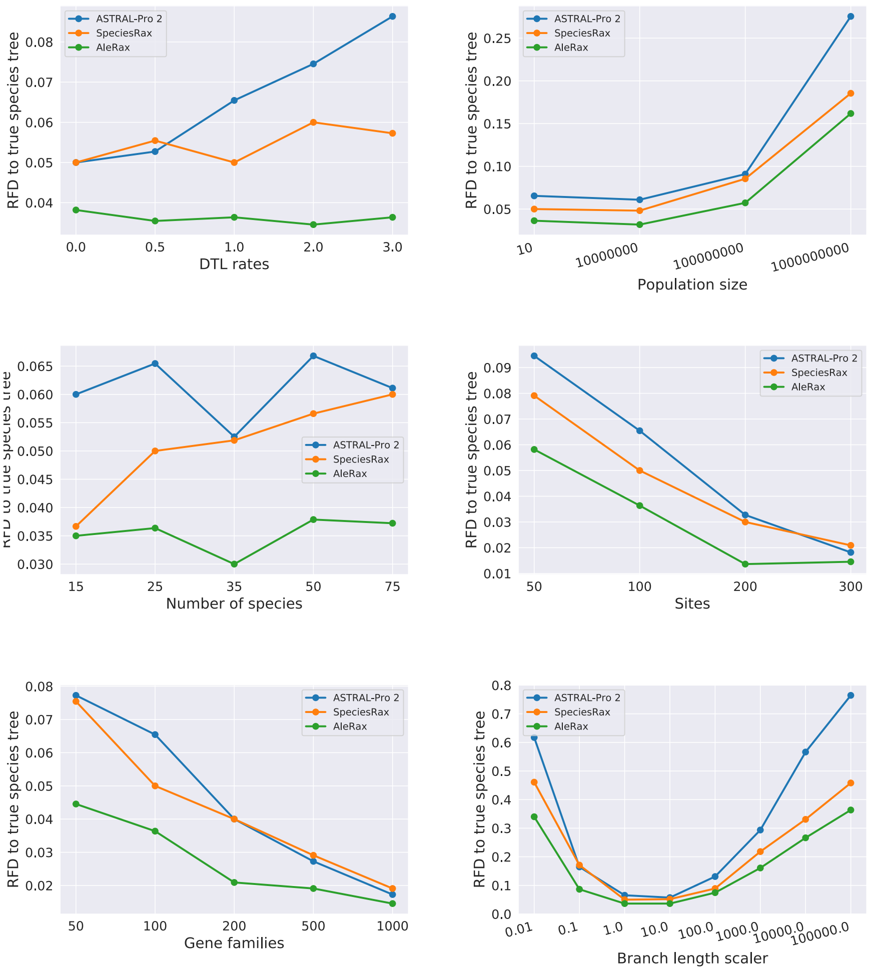


Figure 2: Average unrooted RF distance between inferred and true species trees, in the presence of duplication and loss (no HGT).

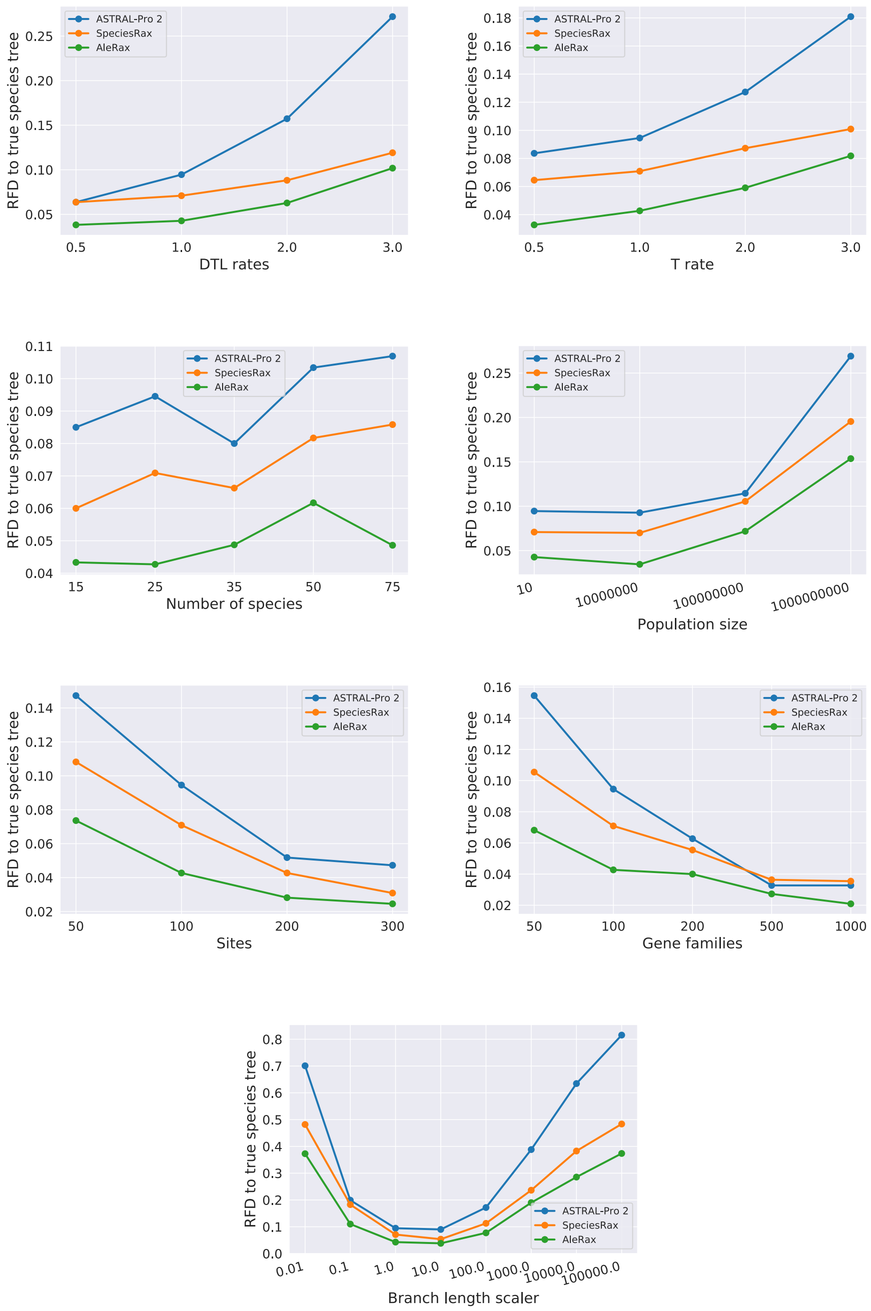


Figure 3: Average unrooted RF distance between inferred and true species trees, in the presence of duplication, loss, and HGT.

| Varying parameters | |
|---|---|
| Dup and loss rate multiplier | 0.5,1.0,2.0,3.0 |
| Number of species | 15, 25, 35, 50, 75 |
| Average number of sites | 50, 100, 200, 300 |
| GFT branch length multiplier | 0.01, 0.1, 1, 10.0, 100.0, 1000.0 |

Table 2: Subset of the DTLSIM dataset used to compare AleRax and ALE on gene tree inference.

#### 4 Benchmarking gene tree inference

In this section, we compare the gene tree accuracy and runtime of AleRax and ALE on a subset of the DTLSIM simulated datasets. We only used a subset of the datasets because of the prohibitive runtimes of ALE on datasets with many species or with very flat gene tree distributions. We list the tested parameter range in Table 2.

We ran ALE on each gene family with the command:

```
ALEml_undated species_tree.newick gene_trees.newick.ale
    sample=100 seed=42 separator=_
```

We distributed the (per-family) jobs over 10 cores.

We ran AleRax on each dataset under the per-family rates mode:

```
mpiexec -np 10 alerax -f families.txt -s species_tree.newick
    --species-tree-search SKIP --gene-tree-samples 100
```

We summarize the results in terms of accuracy in Figure 4 and in terms of runtimes in Figure 5. We observe that AleRax and ALE are on par in terms of accuracy, and that AleRax is on average one order of magnitude faster than ALE.

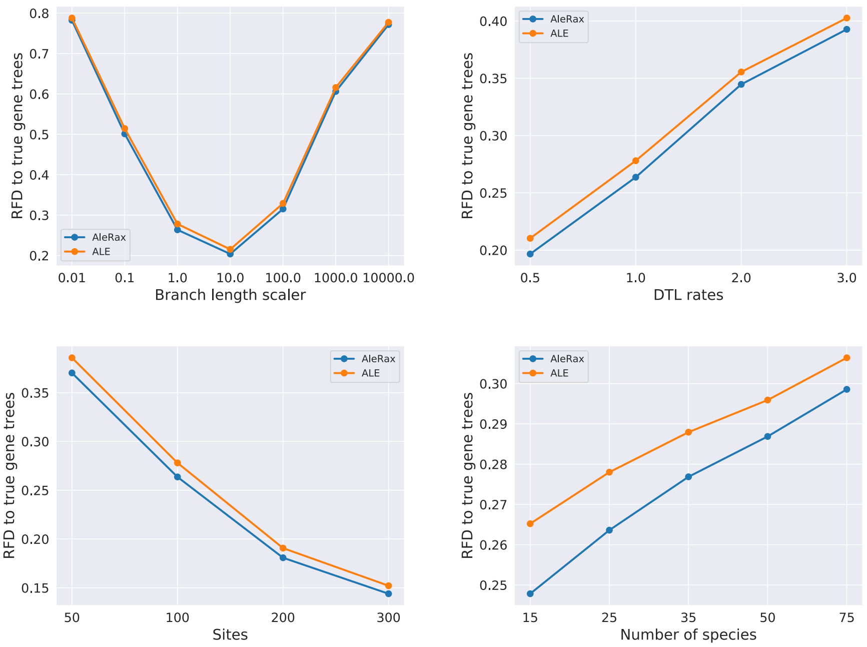


Figure 4: Average unrooted RF distance between sampled and true gene trees on a subset of the DTLSIM simulated datasets.

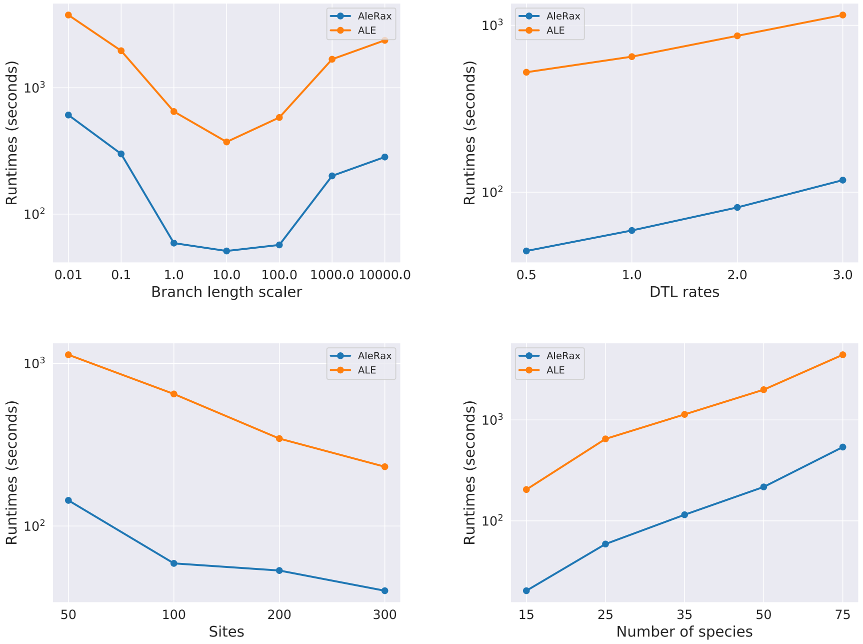


Figure 5: Average runtimes in seconds for reconciled gene tree sampling on a subset of the DTLSIM simulated datasets.

#### 4.1 Benchmarking AleRax vs. ALE on ALESIM

In this section, we describe how we compared the gene tree accuracy of both AleRax and ALE on the ALESIM simulated dataset.

We inferred the distributions of gene trees with MrBayes (Ronquist *et al.*, 2012), with two independent runs, four chains, $1,000,000$ generations, and sampling trees every 1000 generations while discarding the first 100 sampled trees as burnin.

We then ran ALE on each gene family with the command:

```
ALEml_undated species_tree.newick gene_trees.newick.ale
sample=100 seed=42 separator=_
```

We distributed the (per-family) jobs over 30 cores.

We ran AleRax twice, once under the global DTL rates mode ("AleRax"), and once under the per-family rates mode ("AleRax-perfam"):

```
mpiexec -np 30 alerax -f families.txt -s species_tree.newick
    --species-tree-search SKIP --gene-tree-samples 100
```

and

```
mpiexec -np 30 alerax -f families.txt -s species_tree.newick
    --species-tree-search SKIP --gene-tree-samples 100
    --per-family-rates
```

We then computed the average normalized RF distances between the true and the inferred gene trees using the ETE Toolkit (Huerta-Cepas *et al.*, 2016). AleRax-perfam (RF=0.102) and AleRax (RF=0.103) were slightly more accurate than ALE (RF=0.105). We report the results in Figure 6. In terms of runtime, AleRax ($386s$) and AleRax-perfam ($217s$) were one order of magnitude faster than ALE ($3,385s$). Remember that, all those runtimes were obtained by parallelizing the runs over 30 cores, and that AleRax and ALE had a similar parallel efficiency.

#### 4.2 Comparing ALE and AleRax runtimes on the Hogenom database

We extracted $12,408$ gene families corresponding to the 666 core species from the HOGENOM database (Penel *et al.*, 2009). For each gene family, we downloaded both, the maximum likelihood tree, and the set of 1000 ultra-fast bootstrap trees computed with IQTREE-2 (Minh *et al.*, 2020) under the LG+G model. We inferred an unrooted species tree from the maximum likelihood gene trees with Asteroid (Morel *et al.*, 2023) and rooted it manually between Bacteria and Archaea + Eukaryota. We used the ultra-fast bootstrap trees as input gene tree distributions for both ALE and AleRax.

We ran ALE on each gene family with the command:

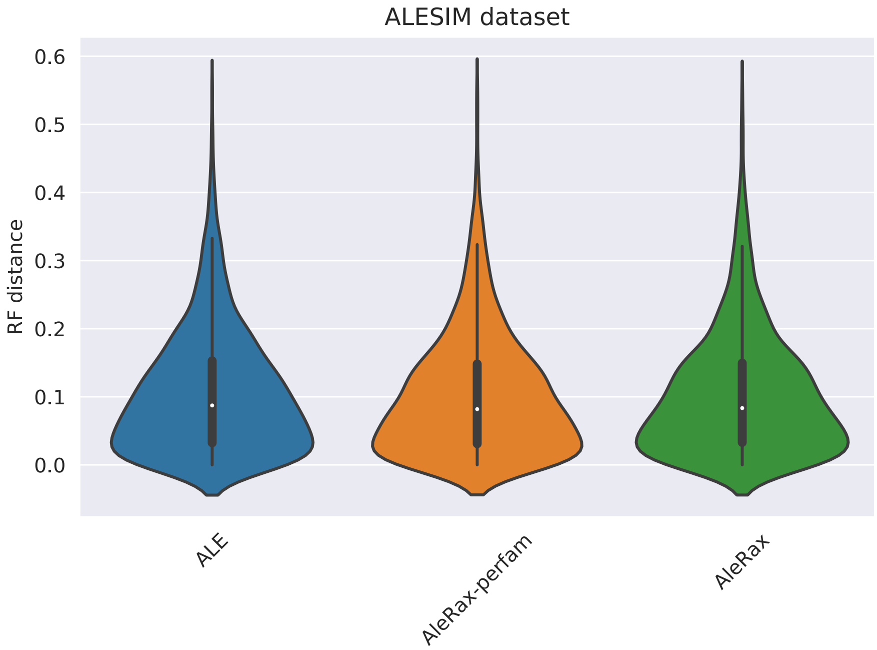


Figure 6: RF distances between the true and the inferred gene trees on the ALESIM dataset.

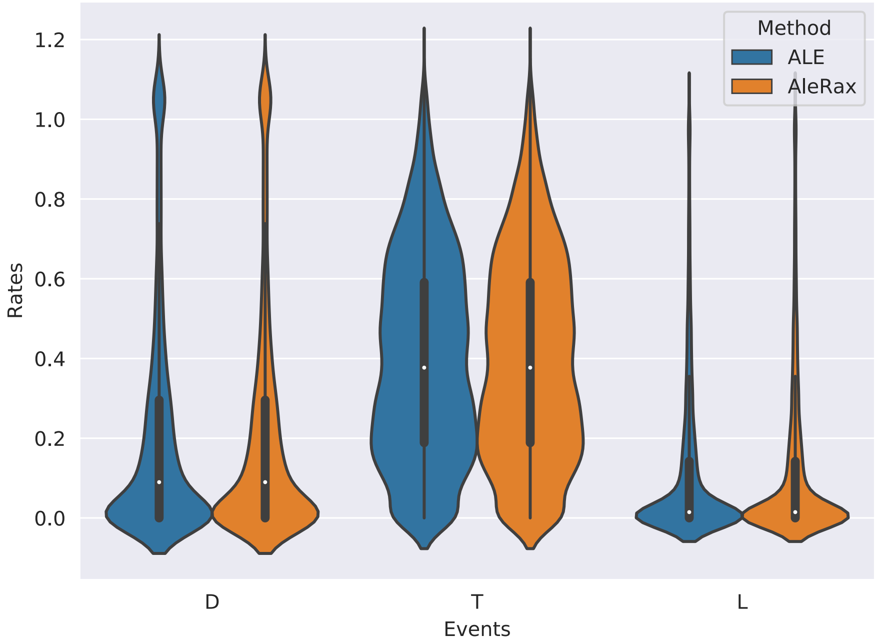


Figure 7: Distributions of the per family DTL rates for both ALE and AleRax runs on the HOGENOM dataset.

```
ALEml_undated species_tree.newick gene_trees.newick.ale
    sample=100 seed=42 separator=_
```

We distributed the (per-family) jobs over 20 cores.

We ran AleRax with per family DTL rates:

```
mpiexec -np 20 alerax -f families.txt -s species_tree.newick
    --species-tree-search SKIP --gene-tree-samples 100
    --per-family-rates
```

AleRax finished after 4.5h and ALE after 44h. AleRax and ALE estimated almost identical distributions of per-family DTL probabilities, as illustrated in Figure 7.
